# Nanoscale imaging of pT217-tau in aged rhesus macaque entorhinal and dorsolateral prefrontal cortex: Evidence of interneuronal trafficking and early-stage neurodegeneration

**DOI:** 10.1101/2023.11.07.566046

**Authors:** Dibyadeep Datta, Isabella Perone, Denethi Wijegunawardana, Feng Liang, Yury M. Morozov, Jon Arellano, Alvaro Duque, Zhongcong Xie, Christopher H. van Dyck, Amy F.T. Arnsten

**Affiliations:** Departments of Neuroscience, Yale University, School of Medicine, 333 Cedar St., New Haven, CT USA 06510; Department of Psychiatry, Yale University, School of Medicine, 333 Cedar St., New Haven, CT USA 06510; Department of Anesthesia, Critical Care and Pain Medicine, Massachusetts General Hospital and Harvard Medical School, 55 Fruit Street, Boston, MA 02114

**Keywords:** pT217-tau, trafficking, plasma, biomarker, entorhinal cortex, dorsolateral prefrontal cortex

## Abstract

**INTRODUCTION:** pT217-tau is a novel fluid-based biomarker that predicts onset of Alzheimer’s disease (AD) symptoms, but little is known about how pT217-tau arises in brain, as soluble pT217-tau is dephosphorylated postmortem in humans.

**METHODS:** We utilized multi-label immunofluorescence and immunoelectron-microscopy to examine the subcellular localization of early-stage pT217-tau in entorhinal and prefrontal cortices of aged macaques with naturally-occurring tau pathology and assayed pT217-tau levels in plasma.

**RESULTS:** pT217-tau was aggregated on microtubules within dendrites exhibiting early signs of degeneration, including autophagic vacuoles. It was also seen trafficking between excitatory neurons within synapses on spines, where it was exposed to the extracellular space, and thus accessible to CSF/blood. Plasma pT217-tau levels increased across the age-span and thus can serve as a biomarker in macaques.

**DISCUSSION:** These data help to explain why pT217-tau predicts degeneration in AD and how it gains access to CSF and plasma to serve as a fluid biomarker.

## 1. Background

Recent advances in Alzheimer’s disease (AD) research have revealed a novel fluid-based biomarker, tau phosphorylated at threonine-217 (pT217-tau), in cerebrospinal fluid (CSF) and blood plasma, that reliably discriminates AD from other neurodegenerative diseases [1]. pT217-tau appears in the earliest presymptomatic stages of AD [1, 2] and predicts subsequent cognitive decline [3]. Moreover, plasma pT217-tau—among multiple tau species and other biomarkers—has demonstrated the highest accuracy to predict the presence of AD neuropathology, including aggregated tau pathology [4]. Novel diagnostic biomarkers will be critical for assessing efficacy of novel therapeutic treatments aimed at early-stage disease [5, 6]. However, it is unknown why this specific phosphorylated tau species is so effective in predicting brain pathology, as little is known about early, soluble pT217-tau expression in brain. It is especially challenging to address this issue in human brain, as soluble phosphorylated tau is rapidly dephosphorylated *postmortem*, and PET scans detect late-stage, fibrillated tau but not the earlier-stage, soluble pT217-tau that likely gives rise to the signal in CSF and plasma [7]. However, the etiology of pT217-tau in aging brain can be addressed in rhesus macaques, where perfusion fixation allows capture of phosphorylated proteins in their native state [8].

Aging rhesus macaques naturally develop tau pathology with the same qualitative pattern and sequence as humans, including initial cortical pathology in layer II of entorhinal cortex (ERC), and later in layer III of dorsolateral prefrontal cortex (dlPFC) [8]. As with humans, tau pathology begins in dendrites of excitatory neurons, where tau hyperphosphorylation leads to aggregation and subsequent fibrillation, forming neurofibrillary tangles within neurons that die via autophagic degeneration [9, 10]. Neuropathological studies show that tau pathology begins about a decade prior to formation of Aβ plaques [11], and correlates with gray matter loss [12] and cognitive impairment [13]. Recent evidence indicates that phosphorylated tau can spread by “seeding” between interconnected neurons, with the ERC a likely origin early in disease [10, 14-17], a finding that has been replicated in rodent AD models [17-25]. However, little is known about the etiology of pT217-tau in human brain, as both *postmortem* and PET imaging studies can only capture fibrillated tau, which occurs at a much later stage. Perfusion fixation of monkey tissue preserves not only phosphorylation state, but internal membranes and organelle architecture, and thus allows high-resolution imaging of molecular location and interactions with cellular organelles at the ultrastructural level, including the detection of large, autophagic vacuoles as early indicators of neurodegeneration.

The current study examined the expression pattern of pT217-tau in aged macaque ERC and dlPFC, including whether pT217-tau could be seen within neurons with early signs of autophagic degeneration, consistent with its predicting future neurodegeneration and disease. We also used high magnification immunoEM to determine whether there was evidence of pT217-tau trafficking between neurons to “seed” tau pathology in higher brain circuits, interfacing with the extracellular space to become accessible to CSF/blood as a fluid biomarker. As immunoEM requires in depth focus, we examined two cortical areas vulnerable to tau pathology: layer II ERC where cortical pathology first begins, and layer III dlPFC, where tau pathology emerges later. We selected age ranges when pT217-tau would be expressed, but still soluble, and thus able to traffic between neurons, and when we first see signs of autophagic degeneration: i.e., 18-24 years for ERC and 26- 31 years for dlPFC. We also examined pT217-tau in plasma from rhesus macaques across the entire extent of the macaque age span (1-34 years), to determine whether it increases with age and can serve as a fluid biomarker in macaques as well as humans. The data revealed that pT217-tau has extensive accumulation within layer II ERC and layer III dlPFC excitatory neurons that express early signs of autophagic degeneration and can be seen trafficking between neurons where it is exposed to the extracellular space, helping to explain why it is an effective, fluid biomarker of early degenerative events in brain.

## 2. Methods

### 2.1 Electron Microscopy

#### 2.1.1 Animals and tissue processing

Three “early” aged (18-24 years) and five “late” aged (26-31 years) rhesus monkeys (*Macaca mulatta*) were utilized in this study and were maintained and euthanized in accordance with the guidelines of the USDA, the Yale University Institutional Animal Care and Use Committee (IACUC), and in accordance with the Weatherall report. As described previously[26-29], primates were deeply anesthetized prior to transcardial perfusion of 100 mM phosphate-buffered saline, followed by 4% paraformaldehyde/0.05% glutaraldehyde in 100 mM phosphate-buffered saline. Following perfusion, a craniotomy was performed, and the entire brain was removed and blocked. The brains were sectioned coronally at 60 μm on a vibratome (Leica) across the entire rostrocaudal extent of the entorhinal cortex (ERC) and dorsolateral prefrontal cortex (dlPFC; Walker’s area 46). The sections were cryoprotected through increasing concentrations of sucrose solution (10%, 20% and 30% each for overnight), cooled rapidly using liquid nitrogen and stored at -80°C. Sections of ERC and dlPFC were processed for immunocytochemistry. To enable penetration of immunoreagents, all sections went through 3 freeze-thaw cycles in liquid nitrogen. Non-specific reactivity was suppressed with 10% normal goat serum (NGS) and 5% bovine serum albumin (BSA), and antibody penetration was enhanced with 0.3% Triton X-100 in 50 mM Tris-buffered saline (TBS).

#### 2.1.2 Histology and Immunoreagents

Previously characterized primary antibodies for pT217-tau raised in rabbit were used, and complexed with species-specific goat secondary antibodies. The following primary antibodies were used: 1) rabbit anti-pT217-tau at 1:200 (cat# AS-54968, Anaspec). The immunogen used KLH conjugated with synthetic peptides corresponding to human tau at phosphorylated threonine 217; 2) rabbit anti-pT217-tau at 1:200 (cat # 44-744, ThermoFisher). The antiserum was produced against a chemically synthesized phosphopeptide derived from the region of human tau that contains threonine 217; 3) mouse anti-EEA1 at 1:250 (cat# 610456, BD Biosciences). Both primary antibodies for pT217-tau produced comparable results. The specificity and selectivity of the primary antibodies have been characterized using immunoblot, immunohistochemistry, immunofluorescence and ELISA approaches in various tissue types in multiple species [30-35]. Normal sera and IgG-free BSA were purchased from Jackson Immunoresearch. All chemicals and supplies for electron microscopy were purchased from Sigma Aldrich and Electron Microscopy Sciences, respectively.

#### 2.1.3 Single pre-embedding peroxidase immunocytochemistry

As described previously[26], the sections were incubated for 72 h at 4 °C with primary antibodies in TBS, and transferred for 2 h at room temperature to species-specific biotinylated Fab’ or F(ab’)_2_ fragments in TBS. In order to reveal immunoperoxidase labeling, sections were incubated with the avidin-biotin peroxidase complex (ABC) (1:300; Vector Laboratories, Burlingame, CA, United States of America) and then visualized in 0.025% Ni-intensified 3,3-diaminobenzidine tetrahydrochloride (DAB; Sigma Aldrich, St. Louis, MO, United States of America) as a chromogen in 100mM PB with the addition of 0.005% hydrogen peroxide for 8 minutes. After the DAB reaction, sections were exposed to osmification (concentration 1%), dehydration through a series of increasing ethanol concentrations (70-100%) and infiltrated with propylene oxide. Tissue sections were counterstained with 1% uranyl acetate in 70% ethanol. Standard epoxy resin embedding followed typical immunoEM procedures followed by polymerization at 60°C for 60 h. Omission of primary antibodies or substitution with non-immune serum resulted in complete lack of immunoperoxidase labeling. Similarly, labeling was nullified when blocking the biotinylated probes with avidin/biotin.

#### 2.1.4 Electron microscopy and data analysis

For single-label immunoperoxidase immunohistochemistry, sections were mounted on microscope slides and ERC layer II and dlPFC layer III was photographed under an Olympus BX51 microscope equipped with a Zeiss AxioCam CCD camera. Zeiss AxioVision imaging software was used for imaging and data acquisition. All sections were processed as previously described and immunoEM imaging was conducted in ERC layer II and dlPFC layer III [26]. Briefly, blocks containing ERC layer II and dlPFC layer III were sampled and mounted onto resin blocks. The specimens were cut into 50 nm sections using an ultramicrotome (Leica) and analyzed under a Talos L120C transmission electron microscope (Thermo Fisher Scientific). Several plastic blocks of each brain were examined using the 4^th^ to 12^th^ surface-most sections of each block (i.e., 200-600 nm), in order to sample the superficial component of sections, avoiding penetration artifacts. Structures were digitally captured at x25,000-x100,000 magnification with a Ceta CMOS camera and individual panels were adjusted for brightness and contrast using Adobe Photoshop and Illustrator CC.2020.01 image editing software (Adobe Systems Inc.). Approximately, 5000 micrographs of selected areas of neuropil with immunopositive profiles were used for analyses with well-defined criteria.

For profile identification, we adopted the criteria summarized in Peters et al. 1991[36]. Dendritic spines in the PFC are typically long and thin, devoid of mitochondria, with the presence of a noticeable postsynaptic density at asymmetric synapses, and often containing an elaborated spine apparatus, the extension of the smooth endoplasmic reticulum into the spine. Dendritic shafts were typically round in perpendicular planes or irregular shaped when assessed in horizontal planes, usually containing mitochondria and numerous tubular and pleomorphic cellular organelles. Depending on the proximity to axon terminals, various dendritic shafts received synaptic inputs. Axon terminals contained accumulations of synaptic vesicles and the axoplasm of these terminals usually contained neurofilaments and mitochondria. The synaptic innervations made by these axon terminals were either asymmetric, containing spherical vesicles, or symmetric, containing pleomorphic vesicles, with typical differences in postsynaptic density, respectively. Unmyelinated axons were small, round processes with a predominantly even and regular shape, traversing the neuropil in a straight orientation, often forming bundles in perpendicular planes. Astrocytic processes were typically of irregular morphology, forming contours that filled the empty space around neuronal elements.

In order to evaluate whether there was concordance between dendrites immunopositive for pT217-tau and signatures of autophagic degeneration, quantitative assessments were performed on a series of low and high magnification 2D micrographs of ERC layer II and supragranular dlPFC layer III. Each field covered an area of 15-30 µm^2^ captured from the thin sections in early aged ERC and late aged dlPFC in rhesus macaques for analyses. Approximately, 200 micrographs of selected areas of neuropil with immunopositive profiles were used for quantitative analyses. The frequency of pT217-tau positive elements was highly consistent across all animals and pooled across animals within an age group.

### 2.2 Immunofluorescence with confocal microscopy

Immunofluorescence staining was carried out on free-floating sections as described previously [37]. Three adult (8-10 years), two “early” aged (18-19 years) and four “late” aged (28-34 years) rhesus monkeys (*Macaca mulatta*) were utilized in this study. Antigen retrieval was performed with 10 mM Solution Citrate Buffer in a hot water bath for 25 min at high temperature. The free-floating sections were left to cool for 30 min at RT. The sections were transferred to 1X TBS for 10 min. Sections were then incubated in 0.5% Sodium Borohydride in 0.01M TBS. Sections were blocked for 1 h at RT in 1X TBS containing 5% bovine serum albumin, 2% Triton X-100, and 10% normal goat serum. Sections were incubated for 72 h at 4°C with pT217-tau primary antibody (dilution 1:100; see supplier information above), MAP2 (dilution 1:1000; ab5392, Abcam) followed by incubation overnight at 4°C with secondary antibodies (1:1000, Alexa-Fluor conjugated, Invitrogen). Following incubation in secondary antibodies, they were incubated in 70% Ethanol with 0.3% Sudan Black B (MP Biomedicals, Cat# 4197-25-5), to decrease autofluorescence from lipofuscin, and counterstained with Hoechst (1:10,000, Thermo Fisher, Cat# H3570). The sections were mounted in ProLong Gold Antifade Mountant (Invitrogen, Cat# P36930). Confocal images from ERC layer II and dlPFC layer III were acquired using a Zeiss LSM 880 Airyscan Confocal Microscope, with the C-Apochromat 40x/1.2 W Korr FCS M27 (Zeiss) water objective. Emission filter bandwidths and sequential scanning acquisition were set up in order to avoid possible spectral overlap between fluorophores. Images were merged employing Fiji software.

### 2.3 Measurement of pT217-tau in blood plasma by Nanoneedles

Fifteen young (1-17 years) and twenty-one aged (18-34 years) rhesus monkeys (*Macaca mulatta*) were utilized in the plasma pT217-tau study (total cohort N=36). We developed in-house phosphorylated tau assays using the nanoneedle technology to measure plasma concentrations of pT217-tau. More than 20,000 nanoneedles are integrated on a silicon substrate assigned to detect one analyte. Each nanoneedle is a single molecule biosensor functionalized with antibodies. Since phosphorylated tau in blood samples are present at low abundance, the binding events of phosphorylated tau on the nanoneedle sensor array follow Poisson statistics, i.e., one or no molecule is captured on each nanoneedle. This allows precise quantitation of analytes by digitally counting the number of nanoneedles that have a positive signal. In the present study, we developed sandwich-type pT217-tau assays to measure the ultra-low concentration of protein levels in patient blood samples. The nanoneedle chip was provided by NanoMosaic (Waltham, MA, USA). Nanoneedles are fabricated in an array format with a spacing of 1.8 μm. Each nanoneedle has a diameter of less than 100 nm. The nanoneedle surfaces were modified with 0.5% 3- aminopropyltrimethoxysilane (APTMS) and activated with 2% glutaraldehyde. This enables antibodies to covalently bind on the surface of the nanoneedles [38]. We used 5 μg/ml capture antibody concentration and incubated with chip overnight. We then washed the chip in PBS and incubated it with blocking solution for 1 hour. 5 μL of plasma samples were thawed and diluted 2 X into the dilution buffer provided by NanoMosaic Inc. The diluted sample was incubated on the chip for 2 hours. After washing, a biotinylated detection antibody at 0.5 μg/ml was incubated on the chip. Next, 0.5 μg/ml streptavidin-HRP conjugate was incubated for 30 minutes, and 3,3’,5,5’- tetramethylbenzidine (TMB) provided by NanoMosaic was applied on the chip for 15 minutes to illuminate the HRP. The enzymatic reaction produced a non-soluble precipitate on the nanoneedles when the sandwich complex was present. The precipitate changes the local refractive index of the nanoneedle and induces a color shift intrinsic to the nanoneedle property. The nanoneedles are imaged under a dark field configuration with a CMOS color camera before and after the assay using the Tessie^TM^ instrument from NanoMosaic. Software provided by NanoMosaic analyzed the colors of all nanoneedles and reported the percentage of the color-shifted number of nanoneedles. We used phospho-specific antibodies to pT217-tau (ThermoFisher 44-744) as the detection antibodies for pT217-tau measurement. The specificity of the phospho-specific antibodies to pT217-tau was validated with peptide array, described in the following section.

The protein levels are reported in relative units specific to the nanoneedle technology (i.e., Nano Unit) as described in previous studies [39, 40] since pT217-tau recombinant protein standards are not available at present. All measurement results were above the lower limit of detection of the nanoneedle assay. Assay specificity was validated with dot blot measurement of the phospho-specific antibodies against an array of synthesized peptides phosphorylated at different residues along the full-length tau protein as described in a previous study [39].

#### Statistical analysis of rhesus macaque blood plasma pT217-tau data

To assess age-related alterations in pT217-tau expression in blood plasma during age-span, a Pearson’s correlation regression analysis was computed. For group analysis, a Welch’s T-test was performed since the pT217-tau expression in blood plasma was normally distributed. Fits were modelled as linear and exponential growth equations.

## 3. Results

### 3.1 Aged rhesus monkeys show progressive development of pT217-tau immunolabeling

Our previous research has demonstrated that aging macaques develop tau pathology in a pattern and sequence qualitatively similar to human AD subjects [8]. Multi-label immunofluorescence labeling for pT217-tau and MAP2, and immunoperoxidase pT217-tau immunolabeling were used to determine the spatial and temporal progression of pT217-tau expression across the age-span in rhesus macaque ERC layer II and dlPFC layer III. In adult (8-10 years) macaque, we observed pT217-tau expression within the nucleus in a subset of excitatory cells **(Figure 1Ai-Ci, Di-Fi)**, which remained in aged animals **(Supplement S1 A-G)**, including delicate expression in proximal apical dendrites and cell soma. Previous studies have identified an important role for tau in preventing genomic instability, rRNA transcription and chromatin relaxation in the nucleus [41-43]. With advancing age, immunofluorescence revealed pT217-tau accumulating in proximal and more distal apical and basal dendrites in ERC layer II excitatory cells and dlPFC deep layer III pyramidal cells, both in “early” aged monkeys (18-19 y; **Figure 1Aii-Cii, Dii-Fii**) and “late” aged monkeys (28- 34y; **Figure 1Aiii-Ciii, Diii-Fiii**).

**Figure 1.**
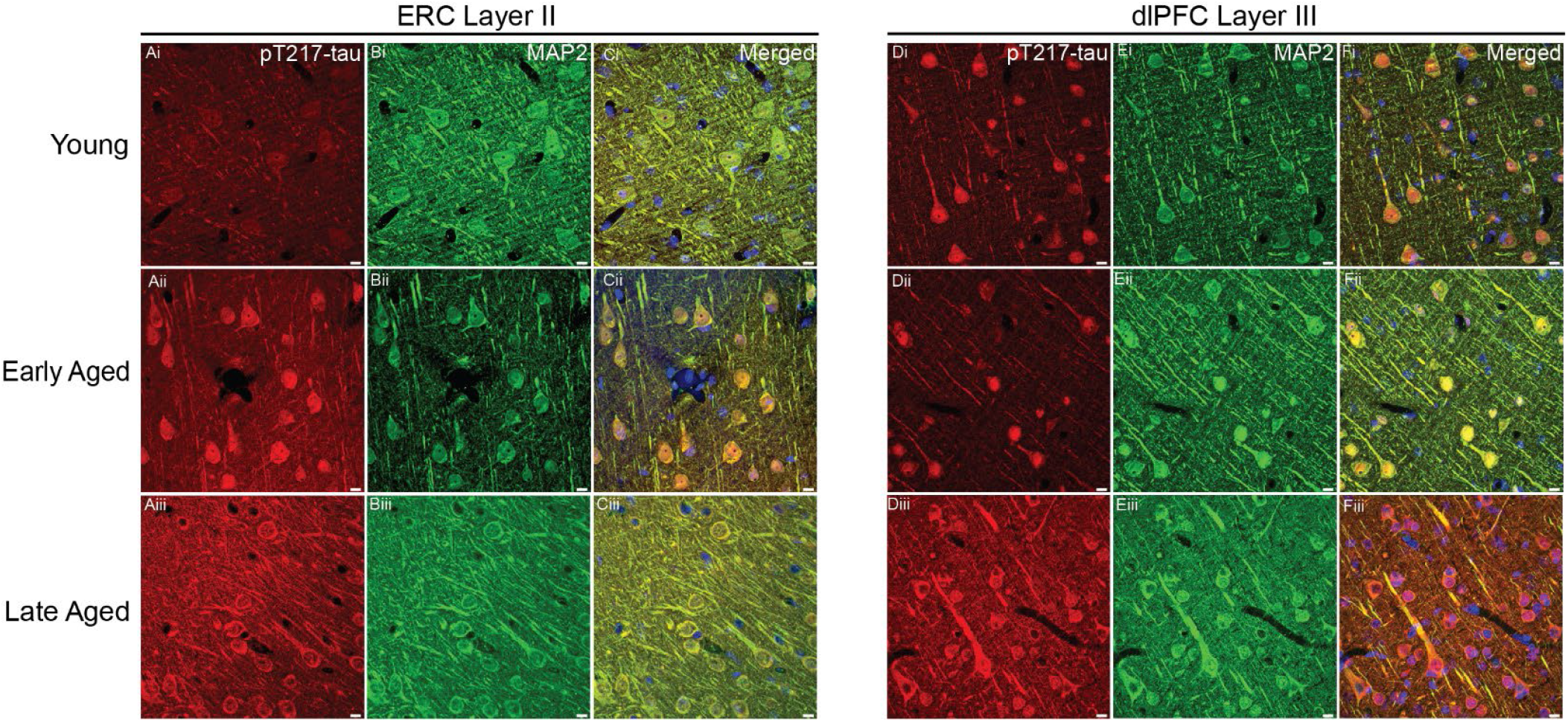
Spatial and temporal localization pattern of pT217-tau across vulnerable cortical regions. Multiple-label immunofluorescence showing pT217-tau labeling (red) co-localized in excitatory neurons (MAP2, green) in ERC layer II and dlPFC layer III across age-span in rhesus macaques. In adult (8-10 years) rhesus macaques, pT217-tau immunolabeling was observed within the perisomatic compartment and the nucleus in a subset of excitatory cells **(Ai-Ci, Di-Fi)**, including delicate expression in proximal apical dendrites in ERC layer II cell islands and dlPFC layer III microcircuits. With advancing age, immunofluorescence revealed pT217-tau accumulating in proximal and more distal apical and basal dendrites in ERC layer II excitatory cells and dlPFC deep layer III pyramidal cells, both in “early” aged monkeys (18-19 y; **Aii-Cii, Dii-Fii**) and “late” aged monkeys (28-34y; **Aiii-Ciii, Diii-Fiii**). Scale bars: 10μm.

### 3.2 Postsynaptic expression of pT217-tau in ERC and dlPFC in aging rhesus macaque

Ultrastructural analyses of “early” aged layer II ERC and “late” aged layer III dlPFC revealed that pT217-tau is concentrated in excitatory neurons in postsynaptic subcompartments within dendritic spines near asymmetric, glutamate-like synapses (**Figure 2**), and in dendritic shafts (**Figure 3**). In dendritic spines, pT217-tau is observed directly in association with the smooth endoplasmic reticulum (SER) spine apparatus near axospinous, presumed glutamatergic, asymmetric synapses, subjacent to the postsynaptic density (PSD) and directly over the postsynaptic membrane **(Figure 2)**. In dendrites, there was extensive pT217-tau aggregation along microtubules and on the SER; these dendrites had the characteristics of excitatory neuron dendritic shafts **(Figure 3)**. The accumulation of pT217-tau along microtubules is particularly relevant to the pathogenesis of AD, as hyperphosphorylation of tau induces detachment from microtubules and primes tau for hyperphosphorylation by GSK3*β*, leading ultimately to the formation of neurofibrillary tangles [44].

**Figure 2.**
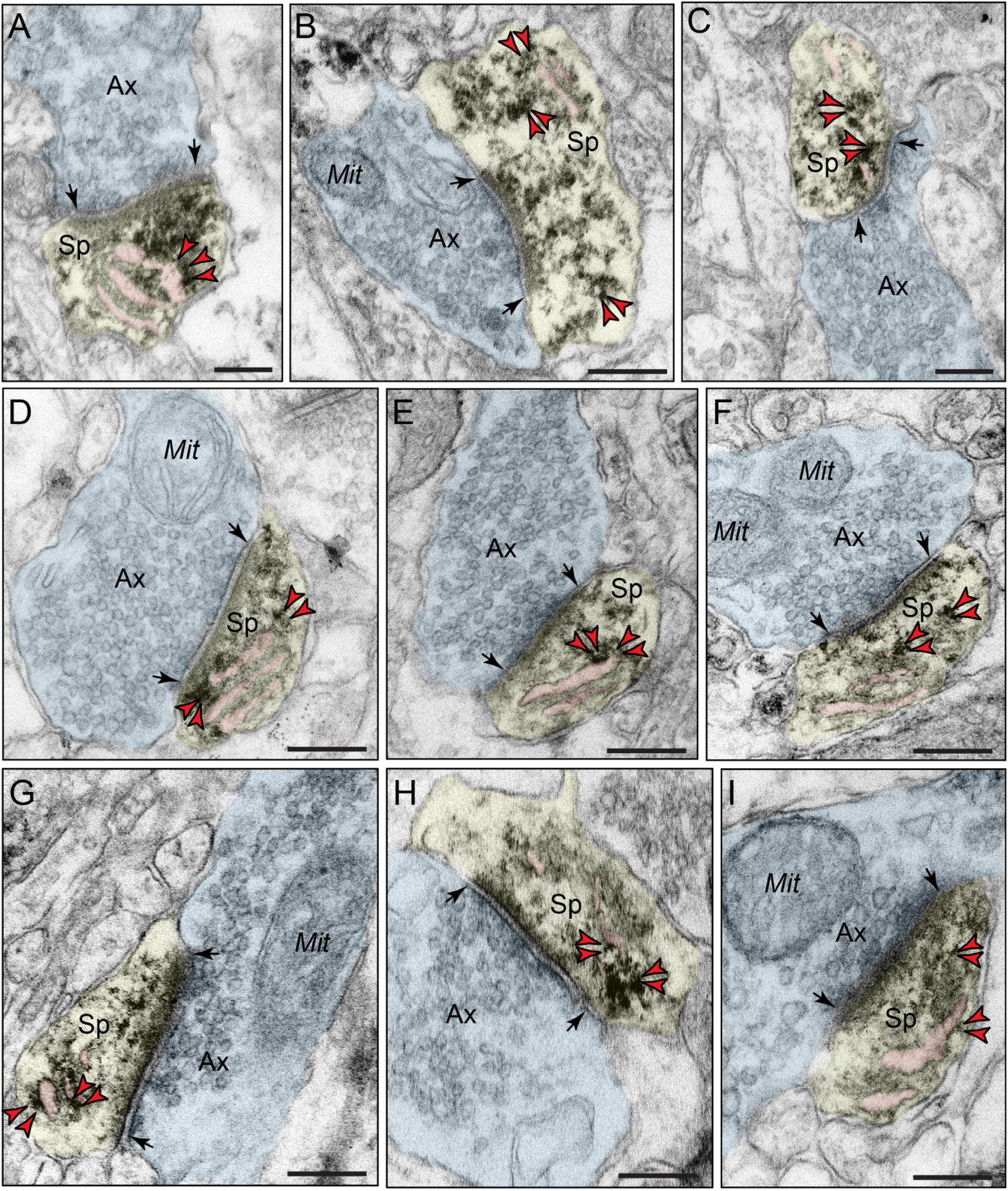
Postsynaptic localization of pT217-tau in dendritic spines in rhesus macaque ERC and dlPFC. Immunoperoxidase labeling revealed that pT217-tau immunolabeling was concentrated in dendritic spines near asymmetric, glutamate-like synapses in rhesus macaque in “early” aged (18-24y) macaque ERC layer II **(A-C)** and “late” aged (26-31y) macaque dlPFC layer III **(D-I)**. pT217-tau was observed directly in association with the smooth endoplasmic reticulum (SER) spine apparatus (pseudocolored pink). pT217-tau immunolabeling is also observed subjacent to the postsynaptic density, near axospinous, asymmetric glutamatergic synapses **(A-C, D, F, G, I)**. Synapses are between arrows. Red *arrowheads* point to pT217-tau immunoreactivity. Profiles are pseudocolored for clarity. Ax, axon; Sp, dendritic spine; Mit, mitochondria. Scale bars, 200 nm.

**Figure 3.**
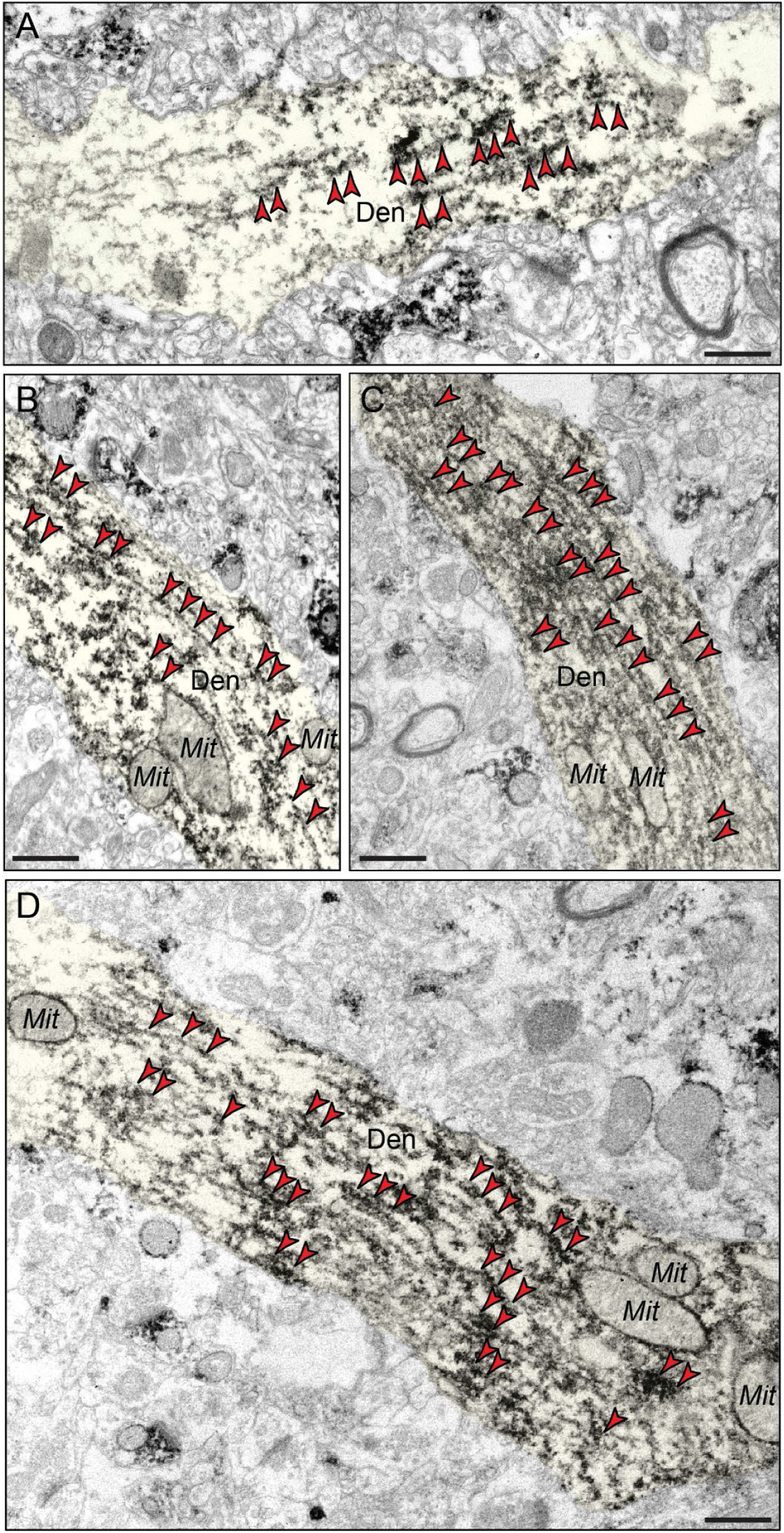
Aggregation of pT217-tau along microtubules in dendritic shafts. ImmunoEM demonstrates aggregation of pT217-tau along dendritic microtubules in putative excitatory cells in “early” aged (18-24y) macaque ERC layer II **(A-B)** and pyramidal cells in “late” aged (26-31y) macaque dlPFC layer III **(C-D)**. Ultrastructural examination revealed dense accumulation of pT217- tau occurred in parallel microtubule bundles in horizontal dendrites. Red *arrowheads* point to pT217-tau immunoreactivity. Profiles are pseudocolored for clarity. Den, dendrite; Mit, mitochondria. Scale bars, 200 nm.

### 3.3 pT217-tau “seeding” within vulnerable neuronal networks in AD

Various lines of evidence in human [10, 14-17] and rodent studies [17-25] have provided evidence of phosphorylated tau trafficking to “seed” pathology in higher cortical networks subserving cognition. ImmunoEM is the only method with sufficient, nanoscale resolution to allow actual visualization of this process *ex vivo*, and thus was of particular interest in the current study. As shown in Figure 4, aggregations of pT217-tau were observed within vesicular structures, including omega bodies fusing with the plasma membrane undergoing endo/exocytosis, within the post-synaptic membrane on dendritic spines in “early” aged (18-24y) macaque ERC layer II **(Figure 4A-B)** and in “late” aged (26-31y) macaque dlPFC layer III **(Figure 4C-D)**. In Figure 4C, pT217-tau could be seen next to both the post-synaptic and pre-synaptic membranes, consistent with pT217- tau trafficking between neurons to “seed” a network of tau pathology. It is noteworthy that pT217- tau was only visualized near excitatory (asymmetric), but not, inhibitory (symmetric) synapses **(Figure 4A-D)**, consistent with tau pathology afflicting glutamatergic, but not GABAergic, cortical neurons in AD [45].

**Figure 4.**
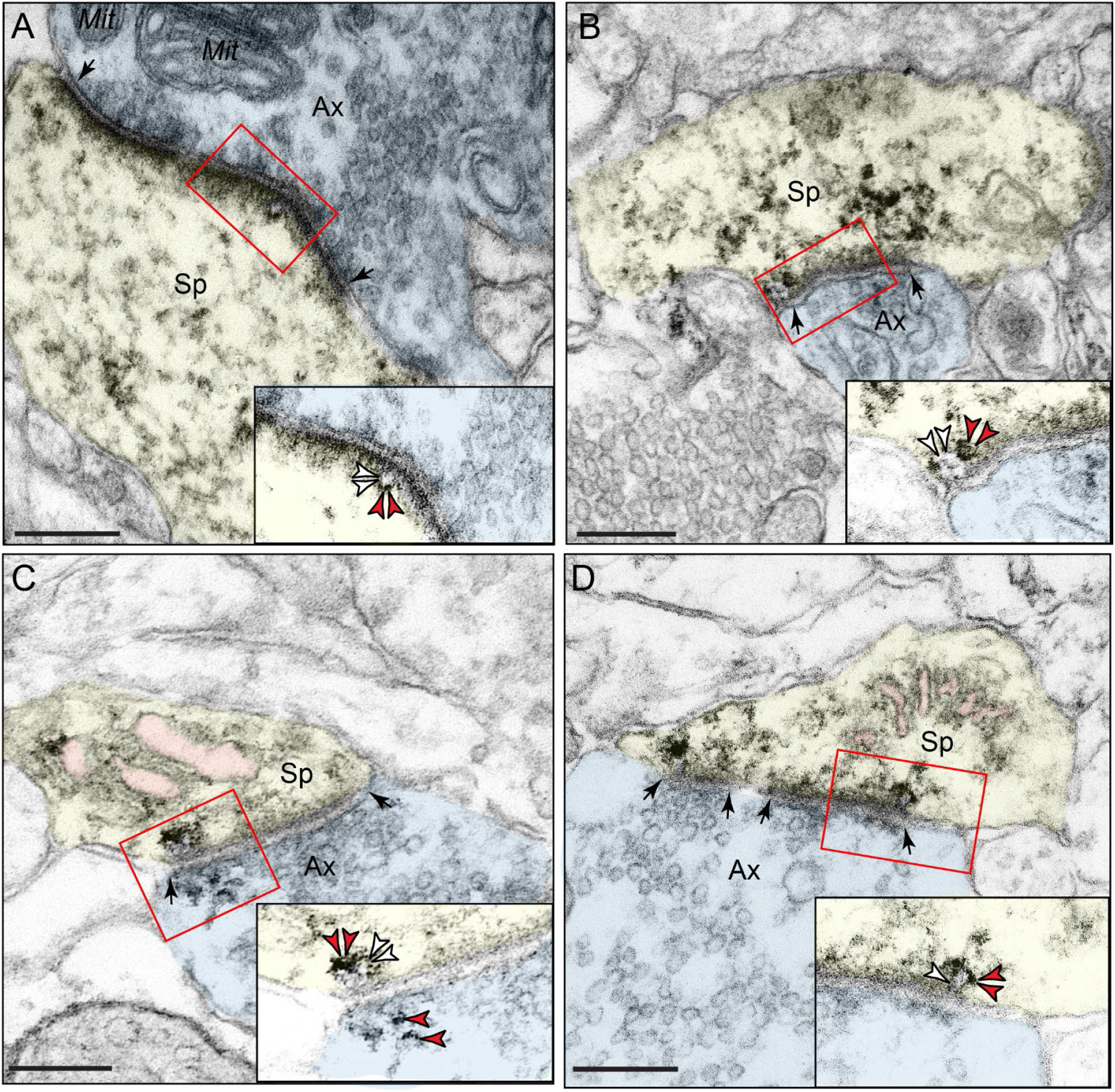
Trans-synaptic trafficking of pT217-tau in aging rhesus macaque. Early-stage, soluble pT217-tau propagates between excitatory neurons within glutamate-like synapses in “early” aged (18-24y) macaque ERC layer II **(A-B)** and in “late” aged (26-31y) macaque dlPFC layer III **(C-D)**. Trans-synaptic propagation of pT217-tau occurs via omega-shaped endosome-like vesicular profiles on the plasma membrane within dendritic spines, specifically within the synapse (insets). All dendritic spines receive axospinous Type I asymmetric glutamatergic-like synapses. Red *arrowheads* point to pT217-tau immunoreactivity. Black *arrows* point to the synapse; white *arrowheads* indicate an omega-shaped profile on the plasma membrane. The SER is pseudocolored in pink. Profiles are pseudocolored for clarity. Ax, axon; Sp, dendritic spine; Mit, mitochondria. Scale bars, 200 nm.

### 3.4 Association of autophagic degeneration in pT217-tau immunopositive dendrites

A critical pathological hallmark in AD involves the formation of autophagic vacuoles, indicative of a degenerative cascade [46-48], oftentimes in NFT-containing neurons, leaving ultimately a “ghost tangle” in advance stages of the illness. Our previous ultrastructural studies have revealed that ERC layer II and dlPFC layer III neurons showed typical signs of AD-like degeneration, including large autophagic vacuoles in the soma and proximal apical and basilar dendrites [8, 49].

We wanted to investigate whether these morphological aberrations are enriched in pT217-tau-immunopositive dendrites. ImmunoEM revealed that pT217-tau is co-localized with autophagic vacuoles in dendrites in “early” aged (18-24y) macaque ERC layer II **(Figure 5A-E)** and in “late” aged (26-31y) macaque dlPFC layer III **(Figure 6A-E)**. Quantitative analyses determined that the vast majority of pT217-tau-expressing dendrites contained autophagic vacuoles. In ERC layer II, we analyzed a total of 116 profiles and found signatures of autophagic degeneration within a large proportion (N=106; 91%) of pT217-tau immunolabeled dendrites **(Figure 5F)**. In dlPFC layer III, we analyzed a total of 82 profiles and found signatures of autophagic degeneration within a large proportion (N=71; 88%) of pT217-tau immunolabeled dendrites **(Figure 6F)**. The robust association between pT217-tau immunopositivity and early signs of autophagic degeneration within dendritic shafts may help to explain why plasma pT217-tau predicts future neurodegeneration and symptoms in humans.

**Figure 5.**
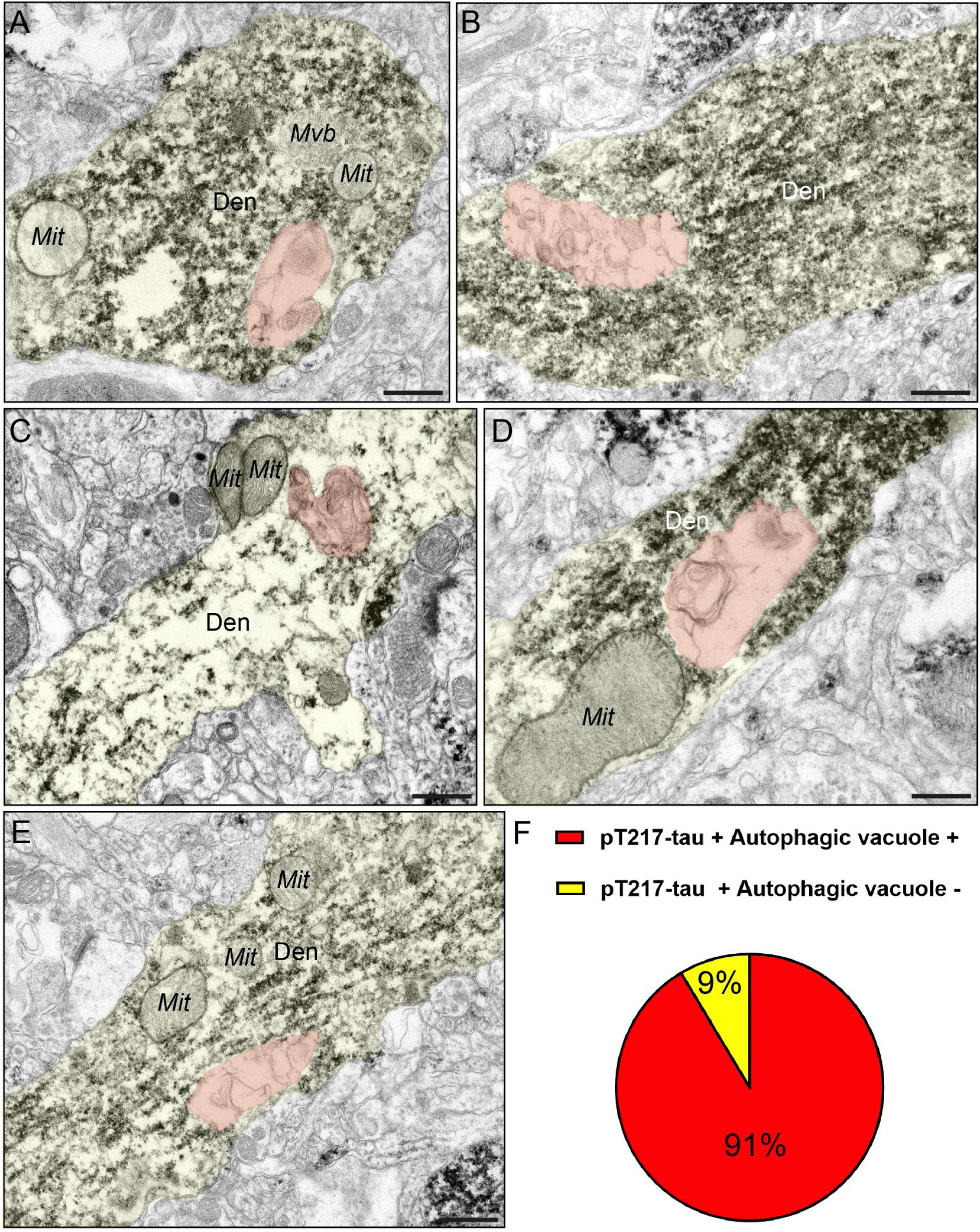
Concomitant autophagic degeneration in pT217-tau immunopositive dendrites in ERC. Association of pT217-tau immunolabeling in degenerating dendrites containing autophagic vacuoles in ERC layer II **(A-E)**. Autophagic vacuoles with multilamellar bodies (pseudocolored in orange) were observed in principal dendrites immunopositive for pT217-tau in “early” aged (18- 24y) rhesus macaque ERC layer II. Systematic quantification (N=116 profiles) in ERC layer II revealed robust concordance between signatures of autophagic degeneration and immunopositivity for pT217-tau (N=106; 91%) within dendrites **(F)**. The percentage of pT217-tau-immunopositive dendrites showing signatures of autophagic degeneration is shown using a pie chart **(F)**. Profiles are pseudocolored for clarity. Den, dendrite; Mit, mitochondria. Scale bars, 200 nm.

**Figure 6.**
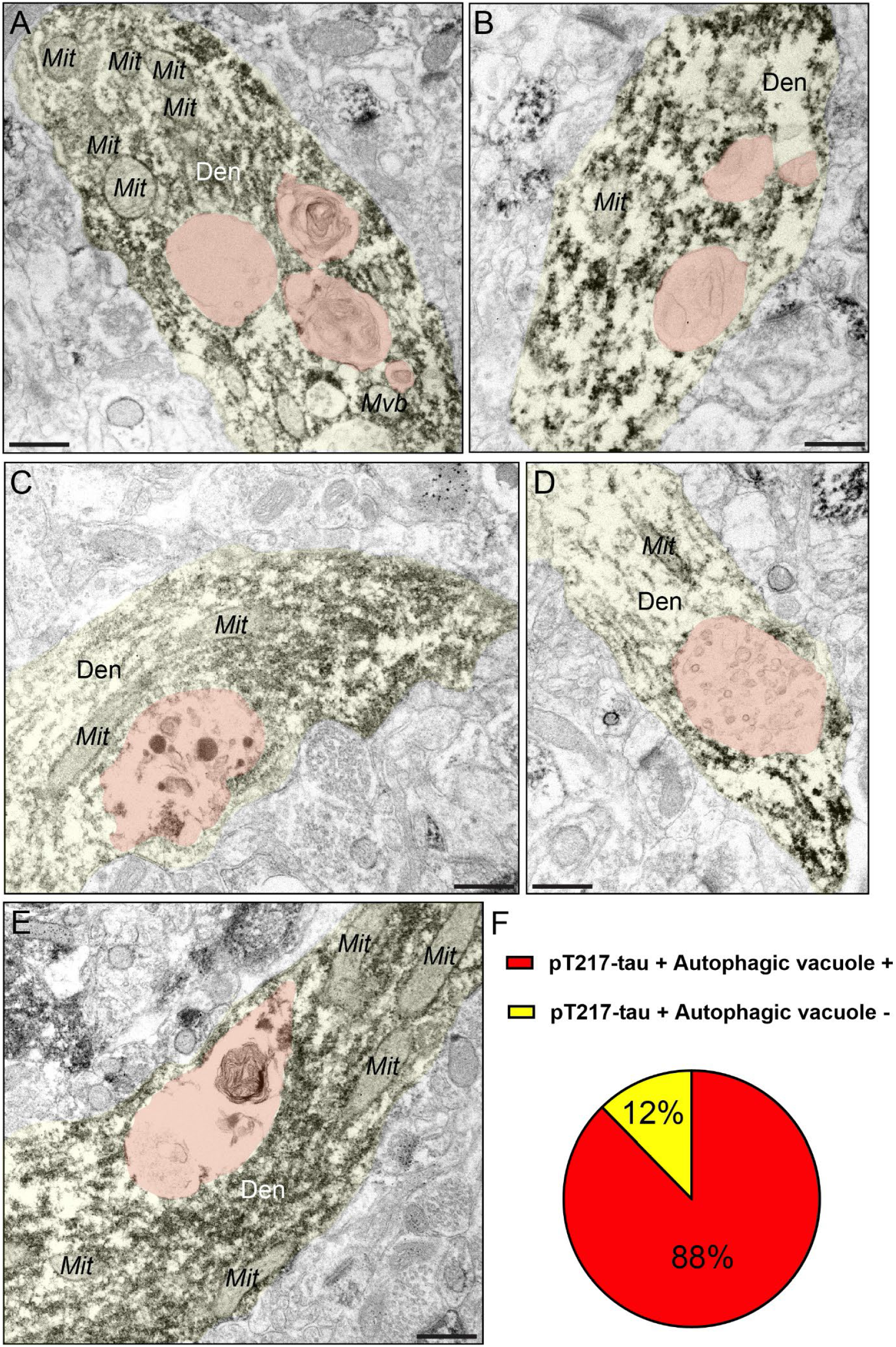
Concomitant autophagic degeneration in pT217-tau immunopositive dendrites in dlPFC. Association of pT217-tau immunolabeling in degenerating dendrites containing autophagic vacuoles in dlPFC layer III **(A-E)**. Autophagic vacuoles with multilamellar bodies (pseudocolored in orange) were observed in principal dendrites immunopositive for pT217-tau in pyramidal cells in “late” aged (26-31y) macaque dlPFC layer III. Systematic quantification (N=82 profiles) in dlPFC layer III revealed robust concordance between signatures of autophagic degeneration and immunopositivity for pT217-tau (N=71; 88%) within dendrites **(F)**. The percentage of pT217-tau-immunopositive dendrites showing signatures of autophagic degeneration is shown using a pie chart **(F)**. Profiles are pseudocolored for clarity. Den, dendrite; Mit, mitochondria. Scale bars, 200 nm.

### 3.5 Aggregated pT217-tau on microtubules “traps” endosomes

Our previous data from aging rhesus monkeys purport a mechanism by which phosphorylated tau may exacerbate A*β* generation within neurons, similar to how genetic predispositions in retromer (e.g., SORL1) signaling pathways may increase the risk of sporadic AD by triggering “endosomal traffic jams,” increasing the time APP spends in endosomes where it is cleaved to A*β* [50]. Endosomes normally traffic on microtubules, providing anterograde and retrograde transport of cargo within cells. In aged macaques, endosomes labeled by EEA1 are often enlarged **(Supplement Fig. S2)**. The current data show that pT217-tau aggregations on dendritic microtubules “trap” endosomes in “early” aged (18-24y) macaque ERC layer II and in “late” aged (26-31y) macaque dlPFC layer III **(Figure 7A-D)**. This disruption of intracellular trafficking may contribute to dendritic pathology and may hasten the cleavage of Aβ from APP (see Discussion). Consistent with dendritic pathology, pT217-tau immunolabeling was also associated with abnormal mitochondria, i.e. mitochondria-on-a-string (MOAS) profiles **(Figure 7A, E-G)**, that have previously been documented in both aging rhesus macaques and patients with AD [51-53].

**Figure 7.**
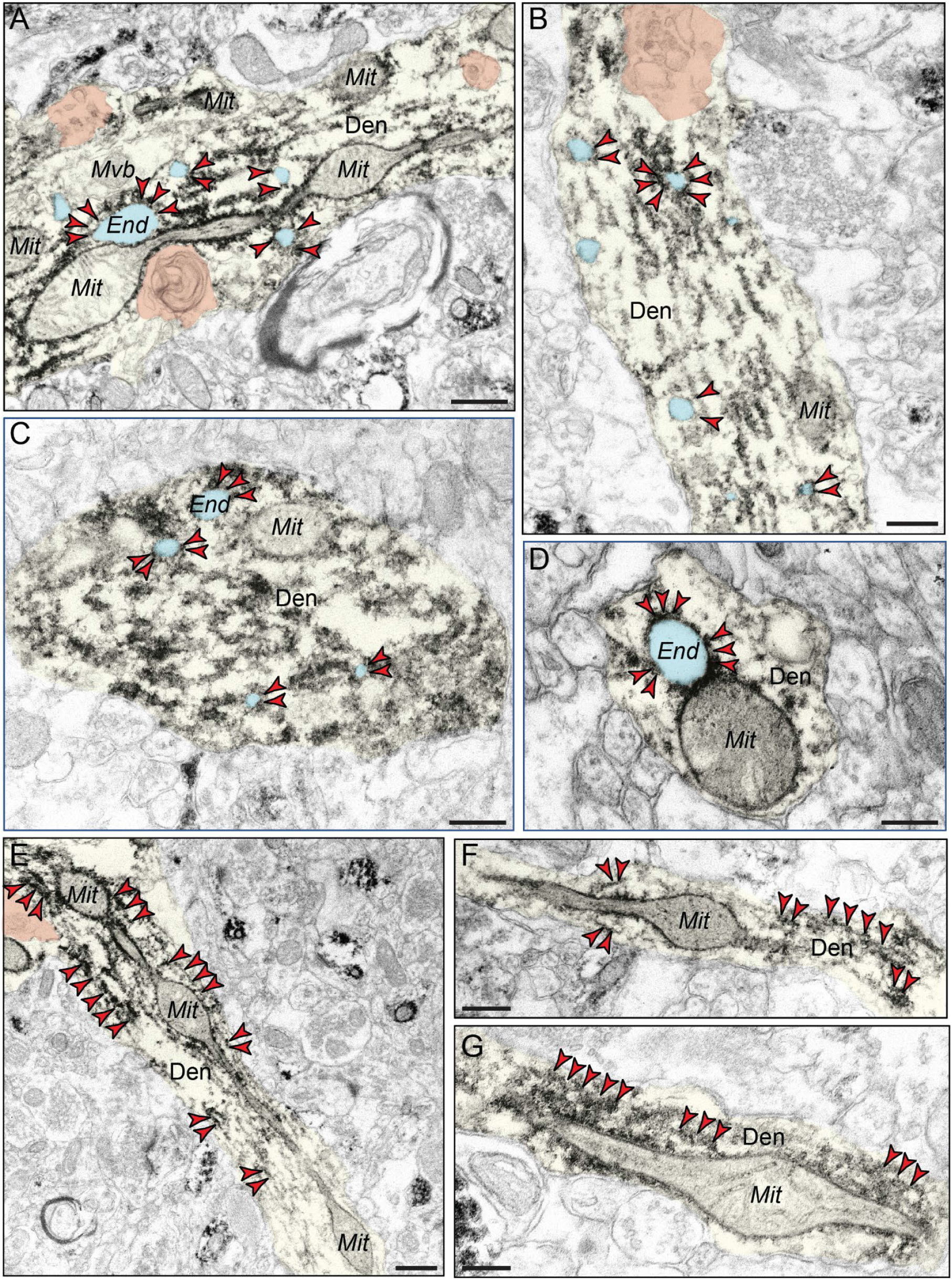
Evidence of pT217-tau trapping endosomes in dendritic shafts, oftentimes near dysmorphic mitochondrial profiles. pT217-tau (indicated by red *arrowheads*) aggregating on microtubules in dendrites where it traps enlarged endosomes (pseudocolored in cyan) in “early” aged (18-24y) macaque ERC layer II and in “late” aged (26-31y) macaque dlPFC layer III **(A-D)**. Aggregations of pT217-tau are often seen near dysmorphic mitochondria, characterized by mitochondria-on-a-string (MOAS) morphological phenotypes, indicative of impaired mitochondrial fission and fusion **(A, E-G)**. MOAS-like morphological disturbances have been described previously in aging rhesus macaques, consistent with local calcium dysregulation. Autophagic vacuolar degeneration (pseudocolored in orange) is also observed in several dendritic shafts, consistent with signatures of neurite dystrophy **(A, B, E)**. Profiles are pseudocolored for clarity. Den, dendrite; Mit, mitochondria. Scale bars, 200 nm.

### 3.6 Age-related increases in pT217-tau in blood plasma in rhesus macaques

Plasma pT217-tau assays are showing promise as indices of brain pathology in humans [1]. The current study examined pT217-tau levels in the plasma of rhesus macaques across the entire macaque age-span. As shown in **Figure 8A**, plasma levels of pT217-tau showed a significant positive correlation with age (R^2^=0.2257; *P*=0.0099) in rhesus macaques. Similarly, a groupwise comparison showed a statistically significant (*P* = 0.0405) increase in pT217-tau in blood plasma in aged (≥18yrs) versus young (<18yrs) animals **(Figure 8B)**. These data suggest that plasma pT217- tau may also be used to track tau pathology in rhesus macaques, where direct comparisons can be made to soluble pT217-tau brain levels, not possible in humans.

**Figure 8.**
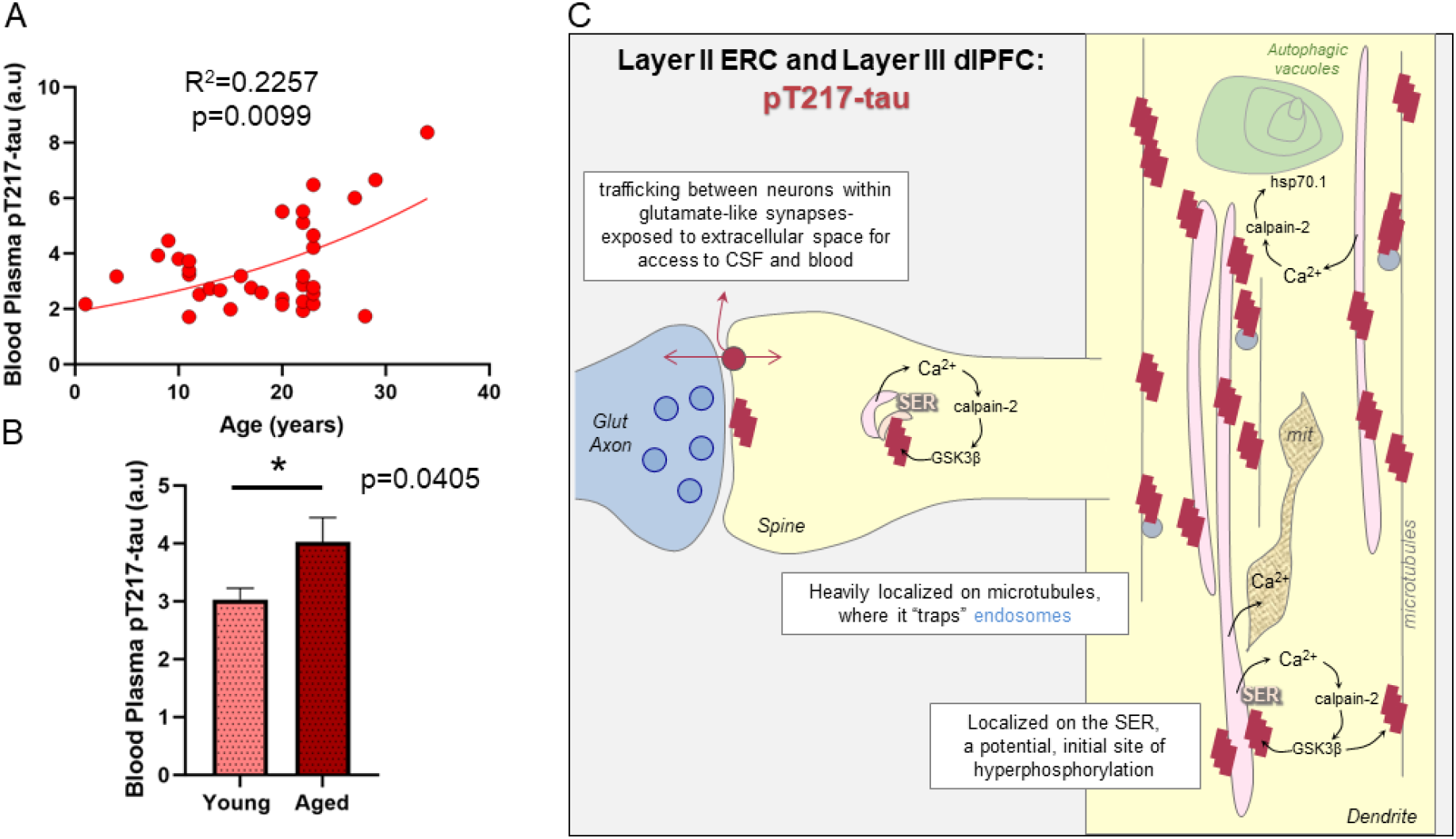
Increased pT217-tau in blood plasma in aging rhesus macaques, and a working model of pT217-tau etiology and toxic actions. Age-related elevation in pT217-tau in blood plasma in aging rhesus macaques **(A)**. Results from a regression analysis in all animals (N=36) was performed and indicated statistically significant increases in pT217-tau signal with advancing age (R^2^=0.2257, p=0.0099). Quantification of pT217-tau in blood plasma grouped by age **(B)**. Young animals (N=15) are compared to aged animals (N=21) via a two-tailed Welch’s t-test (*p=0.0405). SEM is plotted for each group. Summary schematic of pT217-tau expression patterns in “early” aged (18-24y) macaque ERC layer II and in “late” aged (26-31y) macaque dlPFC layer III microcircuits **(C)**. A schematic illustration of pT217-tau localization in dendrites, and its potential etiology and toxic actions. pT217-tau is primarily located in glutamatergic dendrites and dendritic spines, consistent with the known origins of tau pathology in dendrites in humans [45]. The current study found evidence of pT217-tau trafficking between neurons at glutamate-like synapses, interfacing with the extracellular space to become accessible in CSF and plasma. This may contribute to tau “seeding” pathology through an interconnected network of glutamatergic neurons. Aggregations of pT217-tau were prominently expressed on microtubules, as well as on the calcium-storing SER, where increased calcium release may drive GSK3*β* hyperphosphorylation of tau at T217. These dendrites often showed signs of pathology: autophagic vacuoles that are an early sign of autophagic degeneration and abnormal mitochondria (mit), known as “mitochondria-on-a-string”. The aggregations of pT217-tau on microtubules could be seen to “trap” endosomes, interfering with intracellular trafficking needed for healthy dendrites. These “endosomal traffic jams” may also increase the production of A*β* by increasing the time APP spends in endosomes with exposure to *β*- secretase, a hypothesis to be tested in future research.

## 4. Discussion

Our immunoEM and multi-label immunofluorescence analyses of pT217-tau expression in aging rhesus macaque cortical tissue revealed robust expression in the cortical circuits vulnerable in AD. pT217-tau was concentrated in the dendrites and dendritic spines of excitatory neurons in ERC layer II and dlPFC layer III, where it was aggregated on microtubules and on the calcium-storing SER (schematically illustrated in **Fig. 8C**). As calcium dysregulation likely plays an important etiological role in driving tau hyperphosphorylation, the localization on the SER in dendritic spines and dendrites may be related to this mechanism. Particularly relevant to pT217-tau’s role as an emerging fluid biomarker in AD, pT217-tau could be seen trafficking between neurons, where it was exposed to the extracellular space, and thus accessible to CSF and blood (**Fig. 8C**). The association of pT217-tau with dendritic pathology: autophagic vacuolar degeneration, disrupted endosomal trafficking on microtubules, and abnormal mitochondria (**Fig. 8C**), also helps to explain why plasma pT217-tau predicts future neurodegeneration and symptoms in humans. Finally, we found that plasma pT217-tau significantly increases across the age-span in macaques, indicating that this animal model can help to illuminate the relationship between brain and plasma pT217-tau expression, and to test novel therapeutic interventions.

### 4.1 Relevance to etiology of tau pathology in AD

Although postmortem human neuropathological studies have been critical in elucidating the spatial and temporal progression of tau pathology in AD, an important limitation is that these studies are predominantly able to capture fibrillated tau, as biochemical studies of fresh-frozen brain tissue have revealed that soluble phosphorylated tau is rapidly (within minutes) dephosphorylated *postmortem* [54]. In contrast, perfusion-fixation in rhesus macaques captures soluble early phosphorylated tau epitopes *in situ*, and thus allows visualization of the earliest stages of tau pathology that cannot be seen in human. Our data in rhesus macaques show that there is a prolonged stage of early tau pathology accumulation, thus providing an opportunity for timely intervention prior to neurodegeneration. Recent proteomic analyses of post-translational modifications (PTMs) of tau revealed that pT217-tau is a crucial, early phosphorylation epitope, distinguishing AD from other neurodegenerative disorders and indicative of disease progression [55]. Plasma and CSF measures of pT217-tau thus provide a rare window into emerging pathology in human brain that cannot be seen with PET imaging or in *postmortem* human tissue, which predominantly capture late-stage, fibrillated forms of tau [56-59]. Our results reveal an age-related increase in plasma pT217-tau in rhesus macaques, suggesting that plasma pT217-tau can be used as an index of ensuing pathology in rhesus macaques as well, where future studies can directly examine how plasma levels relate to brain measures. They may also provide the opportunity to test novel therapeutic strategies without sacrificing invaluable aged macaques.

The current data are also consistent with emerging data showing that calcium dysregulation with increasing age and/or inflammation drives tau hyperphosphorylation in both animal models and in human patients [60]. pT217-tau was concentrated on the SER in dendritic spines and dendrites, which is the key source of intracellular calcium within neurons (**Fig. 8C**). Abnormal calcium storage and release from the SER is evident in both sporadic and inherited AD [61], where high levels of cytosolic calcium drive calpain-2-mediated disinhibition of GSK3β and cdk5 to hyperphosphorylate tau [21, 62-64], including at pT217-tau [65]. As tau is normally localized on microtubules, these toxic actions in dendrites would hyperphosphorylate tau on microtubules, causing tau to detach and aggregate, as seen in the current study. The data also show that aggregations of pT217-tau along microtubules trap endosomes, interfering with intracellular trafficking needed for healthy dendrites, and which may contribute to amyloid pathology [7] as discussed below.

### 4.2 Visualization of tau seeding and relationship to pT217-tau as a plasma biomarker

The current data help to demonstrate how pT217-tau transfers from within neurons to the extracellular space where it can be conveyed to CSF and plasma for detection as a fluid biomarker. ImmunoEM is the only method with sufficiently high resolution to visualize the pT217-tau trafficking between neurons in its native state. This is likely a momentary event, and thus capturing the event in process is fortuitous and rare. The data reveal that pT217-tau gets exposed to the extracellular space during this process, and thus may be conveyed to CSF and plasma. It is important to note that this is an early event, suggesting that the seeding of tau pathology between neurons likely occurs at an early stage when phosphorylated tau is still soluble. It is also noteworthy that this study, as well as our previous work with pS214-tau [66], have only seen this occur in or near glutamatergic (asymmetric) synapses, but not GABAergic (symmetric) synapses, consistent with tau spreading from the ERC to “infect” interconnected cortical networks [45].

The current data suggest that phosphorylated tau may need to be in a flexible state, e.g., in soluble form, to effectively transfer between neurons. In contrast, fibrillated tau labeled with the AT8 antibody has a markedly different appearance, with long, inflexible fibrils (>200nm) [8] that would not be amenable to trafficking within the smaller (∼100nm) vesicle-like structures observed here. Consistent with our findings, recent studies have revealed evidence of pT217-tau vesicular structures associated within granulovacuolar degeneration bodies (GVB) -large membrane-bound vacuoles containing aggregated proteins including phosphorylated tau [67]. GVBs are correlated with the spatial and temporal propagation of tau pathology [68]. Future studies can delineate whether trans-synaptic trafficking of pT217-tau depends upon low-density lipoprotein receptor-related protein 1 (LRP1) [69] and bridging integrator 1 (BIN1) [70] which have been shown to be important for the inter-neuronal trafficking of tau. Altogether, these data from non-human primate brains indicate how tau pathology can spread over a lifetime, starting early and eventually compromising the integrity of higher cortical networks [7].

### 4.3 Relevance for the prediction of clinical progression and pathology in AD

Recent research has shown that plasma pT217-tau discriminates AD from other neurodegenerative diseases, with high accuracy similar to key CSF-or PET-based measures [1]. In autosomal dominant AD, pT217-tau begins with the initial increases in aggregated amyloid-β as early as two decades before the development of aggregated tau pathology [2] and symptom onset [1, 2]. In sporadic AD, pT217-tau is already associated with fibrillar amyloid deposition in the earliest presymptomatic stages beginning at subthreshold amyloid levels (20 centiloids) (Rissman et al, *Alzheimer’s & Dementia*, In Press). Moreover, in emerging longitudinal studies, plasma pT217- tau has been associated with cognitive decline in patients with preclinical AD [3]. Finally, plasma pT217-tau—among multiple tau species and other biomarkers—has demonstrated the highest accuracy to predict the presence of AD neuropathology, including aggregated tau pathology [4].

The current data help to illuminate why this tau species is so predictive, as it is expressed in dendrites that show signs of early degeneration, including autophagic vacuoles and abnormal mitochondria. These degenerative signs may also be related to calcium dysregulation, e.g., where calpain cleaves heat shock protein Hsp70.1 to increase autophagic neurodegeneration [71] (**Fig. 8C**). In AD, autophagic vacuoles are found in AT8-labeled neurons indicative of a degenerative cascade [46-48], eventually leaving a residual “ghost” tangle [72]. Thus, pT217-tau in plasma may serve as an early index of dendrites initiating the degenerative process [73].

### 4.4 pT217-tau “trapping” endosomes in dendritic shafts: Potential interaction between tau phosphorylation and Aβ

Extensive research has shown that A*β* cleavage from APP is increased when APP is expressed in endosomes, as they often contain beta secretase, the crucial factor for A*β* production [74]. Indeed, the increased risk of AD from genetic alterations in retromer-dependent signaling (e.g., SORL1, VPS26, VPS35) is thought to involve this mechanism, where “endosomal traffic jams” increase APP time spent in endosomes where it is cleaved to A*β* [50]. Our data have suggested that aggregations of phosphorylated tau on microtubules may have a similar effect, trapping endosomes to create a similar “traffic jam” [7], including immunoEM evidence of AT8-labeled phosphorylated tau trapping endosomes labeled with APP [8]. The current data show extensive pT217-tau trapping endosomes, suggesting that pT217-tau may be involved in the etiology of amyloid pathology. This hypothesis may help to explain why plasma pT217-tau levels correlate so highly with amyloid pathology in humans [59, 75]. Although it is widely considered that A*β* drives tau phosphorylation, the current data suggest that the converse may also be true and that the extensive aggregation of pT217-tau on microtubules may help to explain the correlation with this particular p-tau species.

In closing, these studies in aged monkeys help to reveal the cell biological processes associated with pT217-tau in ways not possible in human tissue. The data are consistent with pT217-tau having access to the extracellular space to serve as a fluid biomarker that predicts emerging degenerative pathology in the circuits most vulnerable in AD.

## Acknowledgements

We thank Lisa Ciavarella, Sam Johnson, Tracy Sadlon and Michelle Wilson for their invaluable technical assistance. We are grateful to Heiko Braak and Kelly del Tredici for their constructive critiques and thoughtful suggestions. The authors also would like to thank the Center for Cellular and Molecular Imaging Microscopy Facility at Yale Medical School for assistance with the work presented here.

## Author Contributions

DD designed and performed the experiments in immunohistochemistry and immunoEM. DD and DW collected and analyzed immunoEM and immunohistochemistry experiments. IP and DD collected and analyzed multi-label immunofluorescence experiments. FL and ZX conducted plasma experiments. YMM, JA, and AD, contributed to the experimental design and provided technical expertise. CHvD provided invaluable material resources and revised the manuscript. AFTA and DD designed the experiments, supervised the study, and wrote the manuscript. All authors read and approved the final manuscript.

## Consent Statement

Consent was not necessary as no human subjects were involved in any studies.

---

**Supplementary S1.**
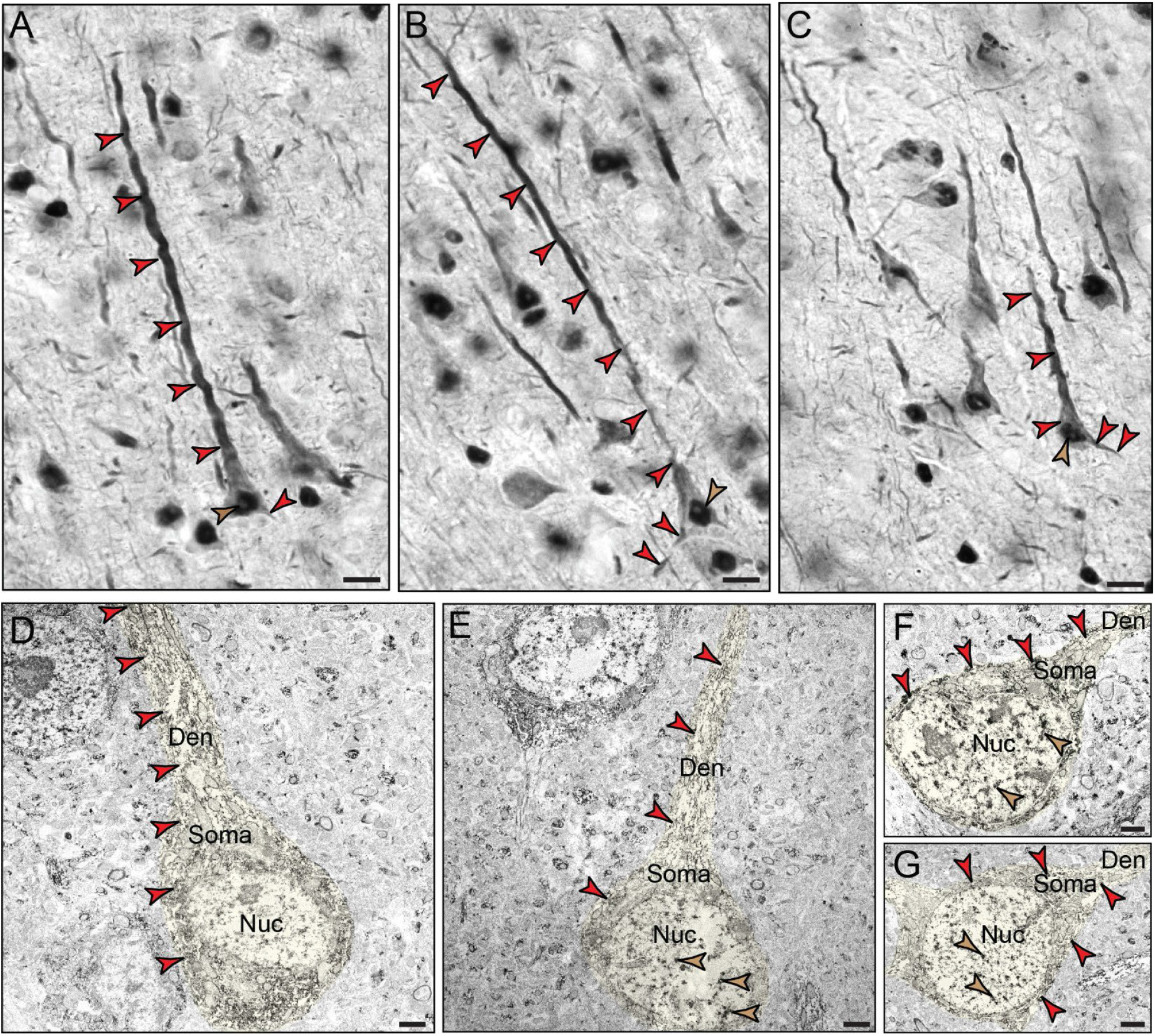
Immunolabeling for pT217-tau in aging rhesus macaques. High-magnification brightfield micrographs revealing intense pT217-tau immunoreactivity along apical and basal pyramidal dendrites (red *arrowheads*), cell soma, and nucleus in aged (30-31 y) rhesus macaque dlPFC layer III **(A-C)**. pT217-tau immunolabeling of pyramidal cells showed aggregated, filamentous pattern within apical dendrites, often with a twisted morphology. Diffuse immunoreactivity is observed in the neuropil, including punctate labeling. Scale bars: 10µm. Low magnification immunoEM micrograph showing pT217-tau immunoperoxidase immunolabeling in pyramidal neurons with a triangular shaped cell body in dlPFC deep layer III **(D-G)**. The pT217-tau immunolabeling is observed in the cell soma and extending along the apical and basilar dendrite (pseudocolored in yellow). Immunolabeling for pT217-tau is also observed in a subset of neurons in the nucleus (indicated by brown *arrowheads*), correlating with light-level immunoperoxidase immunolabeling. pT217-tau protein is also observed in the neuropil. Scale bar: 2µm.

**Supplementary S2.**
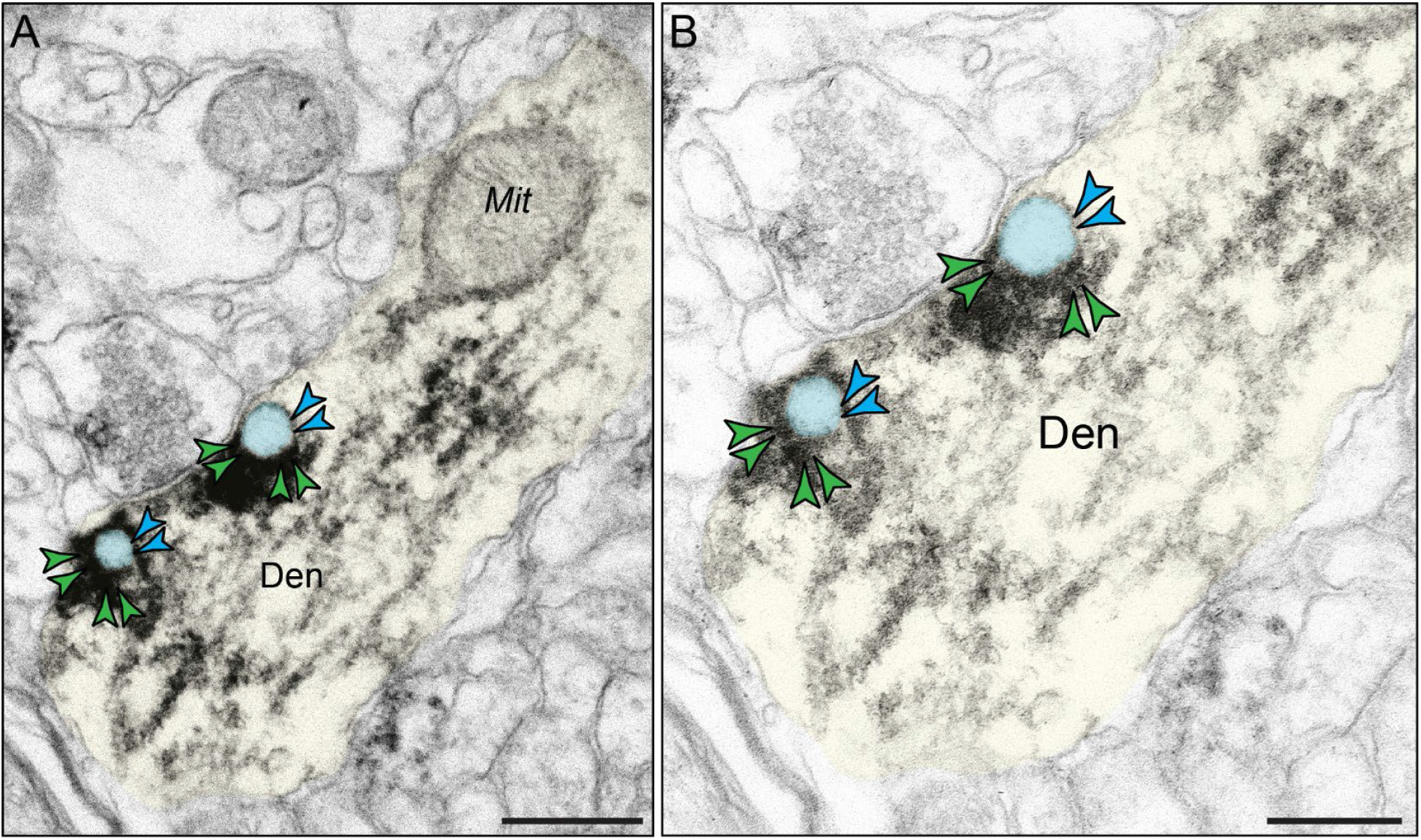
Immunolabeling for endosomes by the marker EEA1 in aged rhesus macaque. ImmunoEM characterization of endosomes (pseudocolored in cyan and cyan *arrowheads*) in aged macaque (25 years) dlPFC layer III using EEA1 (green *arrowheads*), a canonical marker of early endosomes. EEA1 selectively labels endosomes at the ultrastructural level, revealing membrane-bound organelles with a translucent cytosol. Endosomes in aged rhesus macaque appear morphologically enlarged within dendritic shafts near microtubule bundles and mitochondria in aged macaque (25 years) dlPFC layer III **(A)**. Higher-magnification micrograph showing selective labeling of EEA1 with endosomes within dendritic shafts in aged macaque (25 years) dlPFC layer III **(B)**. Den, dendrite; Mit, mitochondria. Scale bars, 200 nm (A), 100 nm (B).

